# Liquid-crystal organization of liver tissue

**DOI:** 10.1101/495952

**Authors:** Hernán Morales-Navarrete, Hidenori Nonaka, André Scholich, Fabián Segovia-Miranda, Walter de Back, Kirstin Meyer, Roman L. Bogorad, Victor Koteliansky, Lutz Brusch, Yannis Kalaidzidis, Frank Jülicher, Benjamin M. Friedrich, Marino Zerial

## Abstract

Functional tissue architecture originates by self-assembly of distinct cell types, following tissue-specific rules of cell-cell interactions. In the liver, a structural model of the lobule was pioneered by Elias in 1949. This model, however, is in contrast with the apparent random 3D arrangement of hepatocytes. Since then, no significant progress has been made to derive the organizing principles of liver tissue. To solve this outstanding problem, we computationally reconstructed 3D tissue geometry from microscopy images and analyzed it applying soft-condensed-matter-physics concepts. Surprisingly, analysis of the spatial organization of cell polarity revealed that hepatocytes are not randomly oriented but follow a long-range liquid-crystal order. This does not depend exclusively on hepatocytes receiving instructive signals by endothelial cells as generally assumed, since silencing Integrin-ß1 disrupted both liquid-crystal order and organization of the sinusoidal network. Our results suggest that bi-directional communication between hepatocytes and sinusoids underlies the self-organization of liver tissue.

## Introduction

The liver is the largest metabolic organ of the human body and vital for blood detoxification and metabolism. Its functions depend on a complex tissue architecture. In the lobule, the functional unit of the liver, blood flows through the hepatic sinusoids from the portal vein (PV) and hepatic artery toward the central vein (CV). The hepatocytes take up and metabolize substances transported by the blood and secrete the bile through their apical surface into the bile canaliculi (BC) network, where it flows towards the bile duct near the PV (*1–3*). Hepatocytes are polarized cells with a unique organization of apical and basal plasma membrane on their surface (*1, 4*). In contrast to simple epithelia, where the cells have a single apical surface facing the lumen of organs, hepatocytes exhibit a multipolar organization, i.e. have multiple apical and basal domains (*1, 4, 5*). Such organization allows the hepatocytes to have numerous contacts with the sinusoid and BC networks to maximize exchange of substances.

Although the general organization of the liver into distinct millimeter-sized lobules is quite clear, the micro-anatomy of a single lobule is much less understood. Sinusoidal endothelial cells and hepatocytes form a heterogeneous 3D packing of cells and labyrinths of sinusoids and BC without apparent order (*1, 5*). However, the function of sinusoid and BC networks prompt precise design requirements: each hepatocyte must be in contact with both networks, yet the networks must never intersect. This defines a problem of self-organization to satisfy these competing design requirements. Hans Elias in 1949 pioneered the structural analysis of the mammalian liver tissue (*6–9*). He proposed a structural model whereby the sinusoids are separated from one another by walls of hepatocytes (one-cell-thick), forming a “*continuous system of anastomosing plates, much like the walls separating the rooms within a building*” (*6, 8*). In his idealized model, Elias proposed that the tissue structure is based on hepatic plates built of alternate layers of polyhedral (decahedra and dodecahedra) cells forming a network of BC and traversed by the sinusoids. The model has been a milestone in the field. However, the analysis underlying the structural model was hampered by the difficulties of reconstructing the 3D tissue structure, which at that time relied on stereological analysis of 2D images (*6, 8–10*). Consequently, the limitations in throughput of 3D reconstructions were a major bottleneck for inferring the rules of structure governing liver tissue. Almost 70 years later, we took advantage of developments in tissue clearing, high-performance microscopy imaging, computer-aided image analysis, and 3D tissue reconstruction to revisit the organizational principles of liver tissue.

## 3D segmentation and quantitative analysis of liver architecture

To understand liver architecture, from the lobule down to the sub-cellular level, we built a 3D geometrical digital representation of adult mouse liver from confocal images (Fig. 1) using a multi-resolution approach (*11*). Mouse livers were fixed, sectioned into 100μm serial slices, and immunostained for cell nuclei (DAPI), cell borders (Phalloidin), hepatocyte apical plasma membrane (CD13), and the extracellular matrix (ECM, fibronectin and laminin) to visualize the basal plasma membrane of hepatocytes facing the sinusoidal endothelial cells (Fig. S1). Full slices and selected regions-of-interest comprising a whole liver lobule were imaged at low resolution (Fig. 1A-C) to determine the position of CV and PV as landmarks, which allowed locating and re-imaging individual CV-PV-axes, at high-resolution (Fig. 1D, corresponding to gray box in Fig. 1C). The multi-resolution imaging allowed us to analyze the distribution of apical and basal surfaces of single hepatocytes (Fig. 1F), as well as sinusoids and BC network geometry (Fig. 1E) in relation to the CV-PV axis (quantitative structural parameters in Fig. S2 and S3). Analysis of thousands individual hepatocytes revealed the full complexity of distribution of the apical, lateral and basal surfaces.

**Fig. 1.**
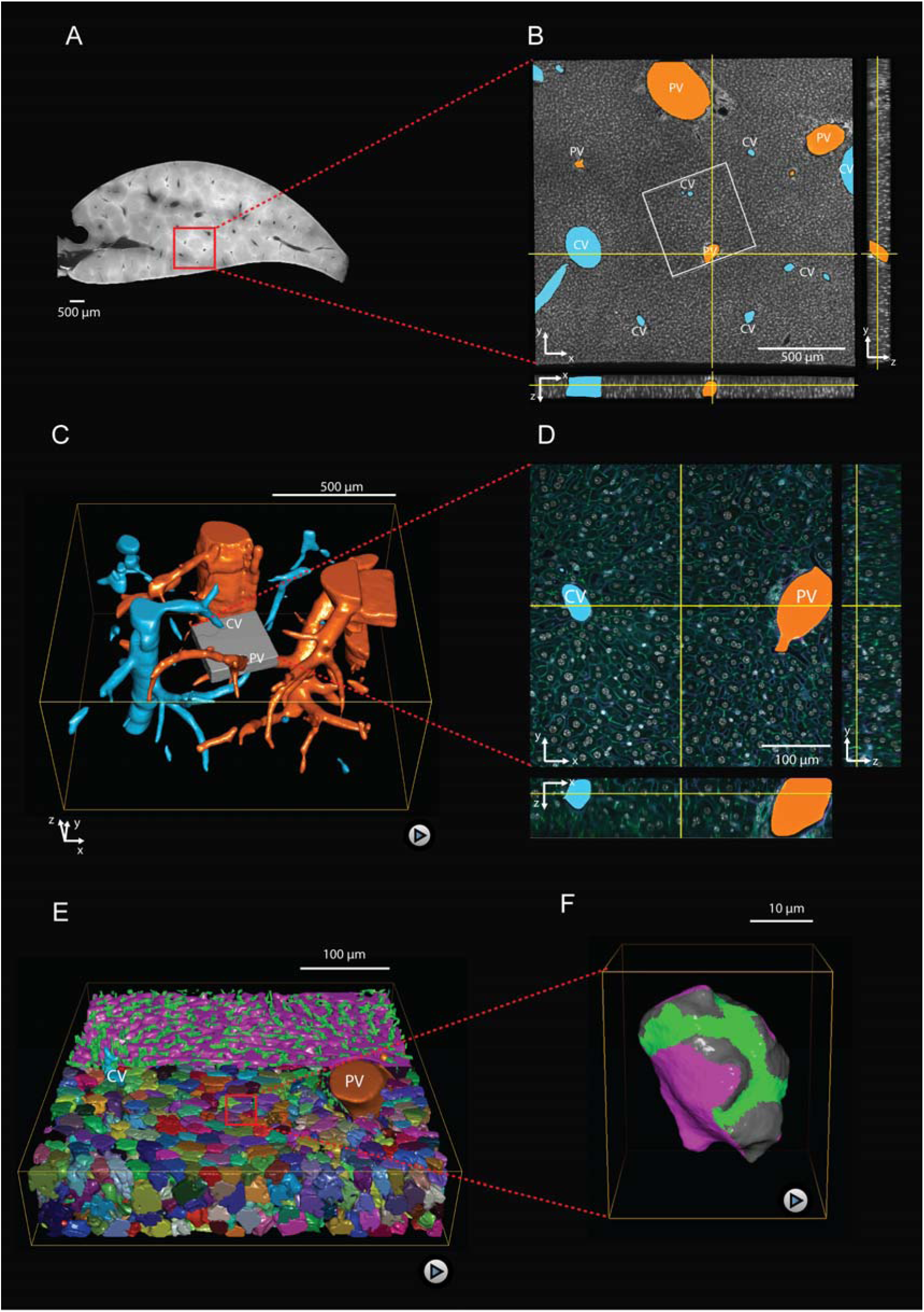
Multi-resolution imaging of the mouse liver lobule. **A, B)** Low-resolution imaging of an optically cleared liver tissue slice, stained for hepatocyte cell borders (cyan, Phalloidin) and nuclei (gray, DAPI); voxel size 1*μ*m × 1μm × 1μm. Central veins (CV, cyan) and portal veins (PV, orange) are highlighted. **C)** 3D reconstruction from a stack of low-resolution images from 10 serial slices. **D)** High-resolution imaging was performed in a sub-region (indicated as gray box in panel C) and stained with 4 different markers for hepatocyte cell borders (cyan, Phalloidin), nuclei (gray, DAPI), hepatocyte apical plasma membrane (green, CD13), and basal plasma membrane (magenta, fibronectin/laminin); voxel size 0.3*μ*m × 0.3μm×0.3μm. **E)** Reconstruction of sinusoidal (magenta) and bile canaliculi (green) networks connecting CV and PV, as well as contacting hepatocytes. **F)** 3D representation of a single hepatocyte showing apical (green), basal (magenta) and lateral (grey) plasma membrane domains.

## Hepatocytes display biaxial cell polarity

The liver parenchyma appears to lack a regular structure at the mesoscopic scale (Fig. 1). Yet, its functional requirements suggest the existence of hidden order. To reveal it, we examined the orientation of hepatocyte polarity in the tissue. In simple epithelia, such as in the kidney and intestine, apico-basal cell polarity can be described by a single vector pointing from the cell center to a single apical pole (*1, 12, 13*) (see schematic in Fig. 2A). However, hepatocyte polarity cannot be described by a single apico-basal polarity axis. Specifically, the spherical harmonic power spectrum of their apical surface patterns peaks not at the first but at the second mode. Therefore, we introduce a new concept of biaxial cell polarity described by nematic tensors (see Methods). Nematic tensors have been used to describe anisotropic structures in physics (e.g. liquid crystals composed of anisotropic units (*14*)) and recently in biology (e.g. 2D epithelial tissues (*15*)). They are characterized by two principal axes (a third axis can be deduced from the other two). The geometric meaning of these axes, here termed the bipolar and the ring axis, is best exemplified in two extreme cases (Fig. 2B, C). In the bipolar case, a marker is concentrated on two opposite poles (Fig. 2B), whereas in the ring case, the marker forms a belt around the cell (Fig. 2C). In the first case, the bipolar axis passes through the two poles (orange axis, *a*_1_), whereas the ring axis is not uniquely defined (it is degenerate in the plane perpendicular to the bipolar axis). In the second case, the ring axis is perpendicular to the plane of the ring and well defined (cyan axis, *a*_2_), whereas the bipolar axis is degenerate. In the case of hepatocytes, the distribution of the apical plasma membrane is in between these two extreme cases, resulting in two well defined perpendicular axes (Fig. 2D). Each of the two cell polarity axes has an associated weight deduced from the nematic tensors (see Materials & Methods), *σ*_1_ for the bipolar axis and *σ*_2_ for the ring axis (Fig. 2E). The distribution of weights is skewed in favor of the belt-like apical surfaces. However, extreme cases described only by a single axis are very rare in the population of hepatocytes. We can define an analogous pair of axes for the distribution of basal plasma membrane, *b*_1_ and *b*_2_, yielding similar results (Fig. S4). Therefore, the polarity of hepatocytes is characterized by two nematic tensors and four axes (*a*_1_, *a*_2_, *b*_1_, *b*_2_). Next, we explored the relationship between apical and basal biaxial cell polarity. We found a preferential parallel alignment for apical and basal axes of different types, *a*_1_ aligned with *b*_2_, and *a*_2_ with *b*_1_ (Fig. 2F). In contrast, the apical and basal axes of the same type (*a*_1_ with *b*_1_ and *a*_2_ with *b*_2_) have preferentially a perpendicular orientation. This anti-correlation corroborates the mutual repulsion between apical and basal surfaces of hepatocytes.

**Fig. 2.**
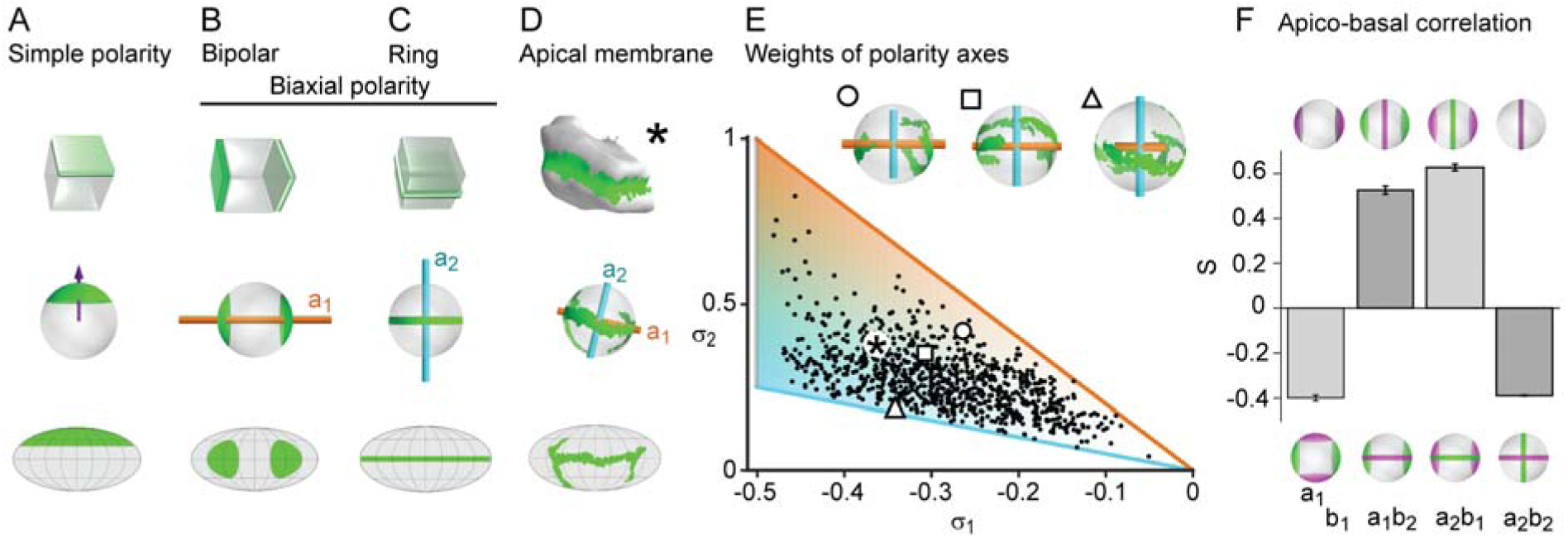
Biaxial cell polarity of hepatocytes. **A)** Idealized representation of simple cell polarity, as found in cells of sheet-like epithelial tissue, showing schematic representation, spherical projection and Mollweide cartographic projection. Simple cell polarity is characterized by a single domain of apical membrane localized at one side of the cell, thus defining a vector (magenta arrow) that points towards the patch of apical plasma membrane. **B, C)** Two extreme cases of biaxial polarity. Biaxial polarity, as introduced here, associates two nematic axes to complex membrane patterns: bipolar and ring axis. We show the bipolar axis (*a*_1_, gold) for the idealized case of two antipodal poles of apical plasma membrane (pure bipolar polarity, ring axis degenerated), and the ring axis (*a*_2_, cyan) for the case of a perfect ring of apical plasma membrane (pure ring polarity, bipolar axis degenerated). **D)** Reconstructed 3D shape of typical hepatocyte with patches of apical plasma membrane (green). The spherical projection is characterized by well-defined bipolar and ring axes. **E)** Respective weights of bipolar axis (σ_1_) and ring axis (σ_2_) for n=857 reconstructed hepatocytes, defined in terms of the eigenvalues of the nematic cell polarity tensor. Extreme cases of pure bipolar or pure ring polarity as shown in B and C correspond to the golden and blue line, respectively. Inset: Spherical projections for three example hepatocytes with corresponding polarity axes (indicated by symbols in scatter plot, corresponding to panel 2A). **F)** Cross-correlation analysis of nematic cell polarity axes for apical and basal plasma membrane patterns reveals that axes of same type are preferentially perpendicular, while axes of different type are preferentially parallel, indicating repulsion between apical and basal plasma membrane domains (n=3 animals).

## Spatial patterns of cell polarity reveal liquid-crystal order in the liver lobule

We next examined the orientation of apical bipolar axes for all hepatocytes within a liver lobule between CV (cyan) and PV (orange) (Fig. 3A). To highlight possible patterns of orientation, we performed local averaging of the bipolar axes over a width of 20*μm*, corresponding to approximately one hepatocyte diameter (Fig. 3B, see Methods). This procedure revealed a spatial organization of hepatocyte polarity with a pattern of the bipolar axis oriented along lines that connect CV and PV. This pattern is reminiscent of flux lines generated by diffusive transport between CV and PV. Indeed, the stationary solution of the diffusion equation with source and sink on CV and PV, respectively, generates a pattern of flux (**J**) (Fig. 3C) similar to the pattern of averaged bipolar apical axes (Fig. 3B). A comparison of averaged apical bipolar axes and the flux pattern **J** is shown in color-code in Fig. 3D and quantified in Fig. 3G, where red color indicates strong alignment with the reference direction and blue denotes a perpendicular orientation. Performing the same analysis for the ring-like axis yielded a preferentially perpendicular orientation with respect to the reference direction **J** (Fig. 3E, G). Consequently, belt-like apical domains are oriented to facilitate bile transport along the reference direction (see also Fig. S5). The simultaneous alignment of two axes, bipolar and ring axis, is indicative of biaxial nematic order(*16*).

**Fig. 3.**
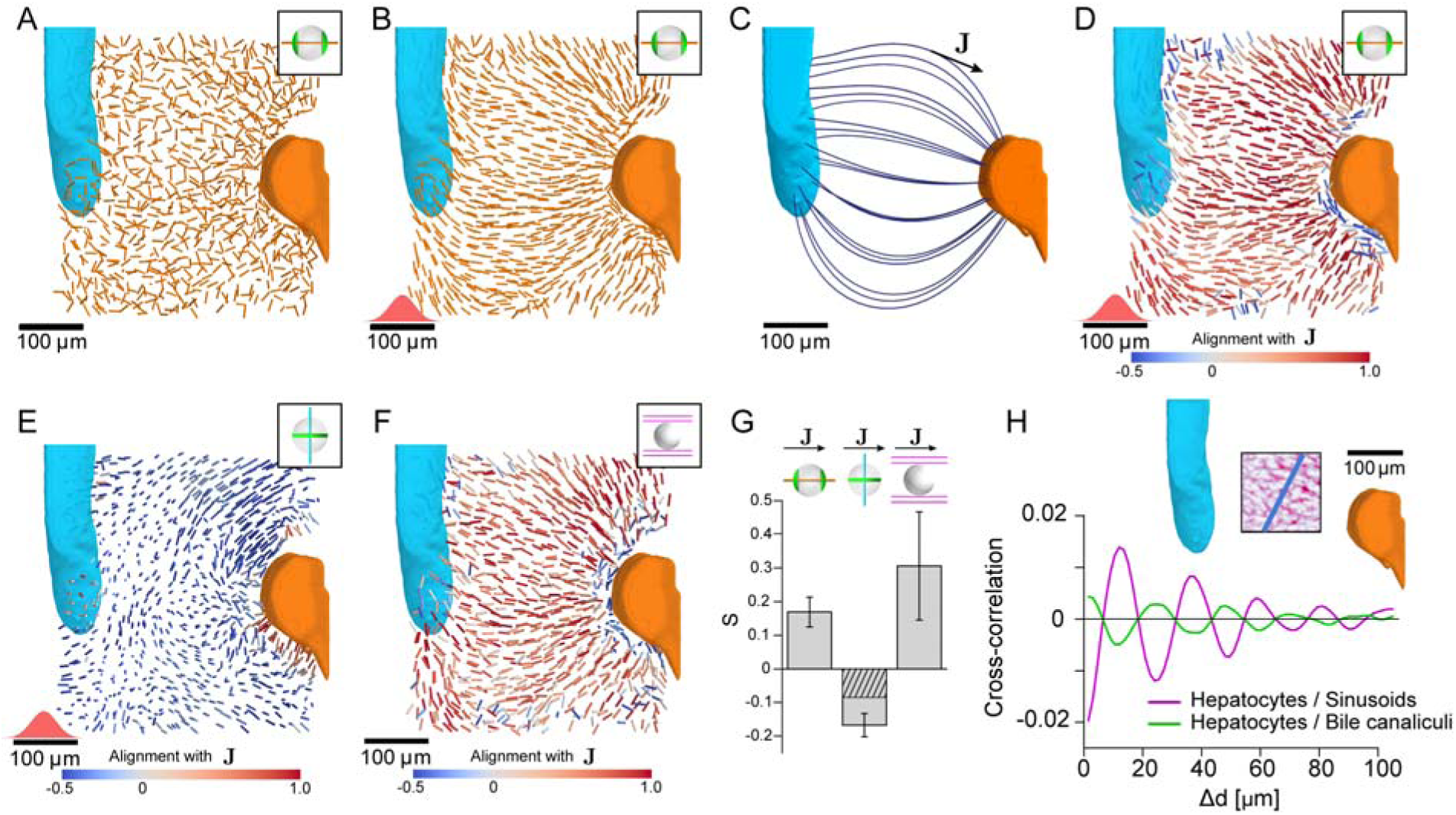
Lobule-level organization of nematic cell polarity. **A)** Bipolar cell polarity axes of apical plasma membrane distribution (*a*_1_) shown as lines of constant length for individual hepatocytes at their respective position in the lobule. **B)** Same as A after local averaging using a 3D Gaussian kernel at each hepatocyte cell center (standard deviation 20*μm*, indicated in red above the scale bar). **C)** Lobule-level reference system with local reference direction (**J**) tangent to flow lines (blue), obtained by solving the diffusion equation with sources and sinks placed at the surface of PV (orange) and CV (cyan). **D)** Same as B, but now axes are color-coded according to their alignment with the local reference direction (**J** defined in panel C. Red colors indicate parallel alignment, whereas blue indicate perpendicular co-orientation. **E)** Same as D, but for the ring axis of apical plasma membrane distribution (*a*_2_). **F)** Same as D, but for the preferred direction of the local sinusoid network surrounding each hepatocyte. **G)** Quantification of alignment with local reference direction for apical bipolar axis, apical ring axis, and preferred sinusoid orientation. The correlation for the apical ring axis exceeds a trivial baseline (hatched bar) that follows from the correlation of the bipolar axes (see Materials and Methods for details) (n=3 animals). **H)** Layered order in the liver lobule. Upper: density of sinusoids in region-of-interest (average density projection along z-axis) and reference direction (blue). Lower: cross-correlation along reference direction between projected density of hepatocytes and sinusoids (magenta) and hepatocytes and bile canaliculi (green). The oscillatory signals reveal layered order with a wavelength of approximately one hepatocyte diameter. All scale bars 100*μm*.

Considering that the sinusoidal network connects PV to CV to ensure blood flow, we expected this network to also exhibit patterns of orientation along the CV-PV axis. To test for this, we determined a local anisotropy axis of the sinusoidal network (see Methods) and found significant parallel alignment between this anisotropy axis and **J**, see Fig. 3F, G. Our analysis shows that cell polarity axes of hepatocytes and the anisotropy of the sinusoid network display biaxial nematic order in the liver lobule. In soft condensed matter physics, such organization is known to result from either weak interactions between anisotropic units, or interactions with an external field, thus creating a liquid crystal-type of organization(*14*). Liquid-crystal order, as found in displays of electronic devices, is characterized by orientational order of basic units, e.g. approximately parallel alignment, yet lack of the translational order of crystalline packings common of solids.

Next, we investigated if translational order could be found at least in one spatial direction. For this, we calculated the cross-correlation for density variations of sinusoids, hepatocytes and BC along a direction perpendicular to the CV-PV axis (see scheme in Fig. 3H). The cross-correlations revealed a periodicity of structures along this direction with a characteristic length-scale of 24*μm*. This periodicity approximately equals the sum of the hepatocyte and sinusoid tube diameters. In contrast, we found no evidence of periodic structures in the direction of the CV-PV axis (see Fig. S6). Such periodicity in only one direction is a hallmark of a layered structure. Therefore, sinusoids, hepatocytes, and BC exhibit a layered organization, with most hepatocytes forming a single layer sandwiched between sinusoidal cells, supporting earlier structural models (*6, 8, 9*).

## Bidirectional communication between hepatocytes and sinusoids is necessary for the maintenance of tissue structure

The layered organization of sinusoids and hepatocytes prompts the question of how coordination between these two cell types is achieved. It has been proposed that the sinusoidal endothelial cells are the main organizers by self-ordering and enforcing the position and, hence, the polarity of hepatocytes (*17, 18*). This requires the establishment of apico-basal polarity with the basal plasma membrane of hepatocytes facing the sinusoidal endothelial cells. A candidate pathway orienting the basal surface of hepatocytes is that of the transmembrane ECM receptors Integrins (*19*). Perturbation of this pathway should result in a flawed coordination between sinusoids and hepatocytes and, consequently, defects in the liquid-crystal order of hepatocyte polarity. Furthermore, if the communication between sinusoids and hepatocytes were unidirectional, the sinusoidal network would be predicted to remain unaltered. To test these predictions, we silenced Integrin-β1 in the liver lobule *in vivo*. The injection of siRNAs formulated into lipid nanoparticles (LNP) provides the advantage of silencing genes with high efficacy and specificity in hepatocytes of adult mice, dosing for either transient or sustained down-regulation (*20, 21*). Injection of siRNAs against Integrin-β1 resulted in a 90% reduction in expression in comparison with control (injected with LNP-formulated Luciferase siRNA), as previously described (21). Loss of Integrin-β1 in liver parenchymal cells led to barely detectable alterations during the first 2–4 weeks. However, after 7 weeks of treatment with Integrin-β1-specific but not control siRNAs, when a significant number of hepatocytes are naturally replaced (22), we detected major alterations in liver tissue organization.

First, the BC network appeared disrupted and more branched (due to an increase of dead-end branches) (Fig. 4A). Interestingly, the biaxial cell polarity of hepatocytes was not compromised. We observed almost unchanged correlation patterns between apical and basal cell polarity axes (Fig. 4B) and indistinguishable distributions of their weights (Fig. S4). This suggests that hepatocyte polarity is maintained by a cell-autonomous mechanism. On the scale of individual hepatocytes, the only change was a significant increase in apical surface at the expense of the basal surface (Fig. S2). However, the long-range order of hepatocyte cell polarity was strongly perturbed (Fig. 4C and Fig. S7). Surprisingly, despite the silencing of Integrin-β1 being limited to hepatocytes at the used dosage, the sinusoidal network was also severely disrupted, with loss of its long-range organization (Fig. 4A, D). This suggests that also hepatocytes provide instructions to sinusoidal endothelial cells. In conclusion, Integrin-β1 KD results in loss of long-range liquid-crystal order of the liver lobule and a perturbed coordination between BC and sinusoidal networks.

**Fig. 4.**
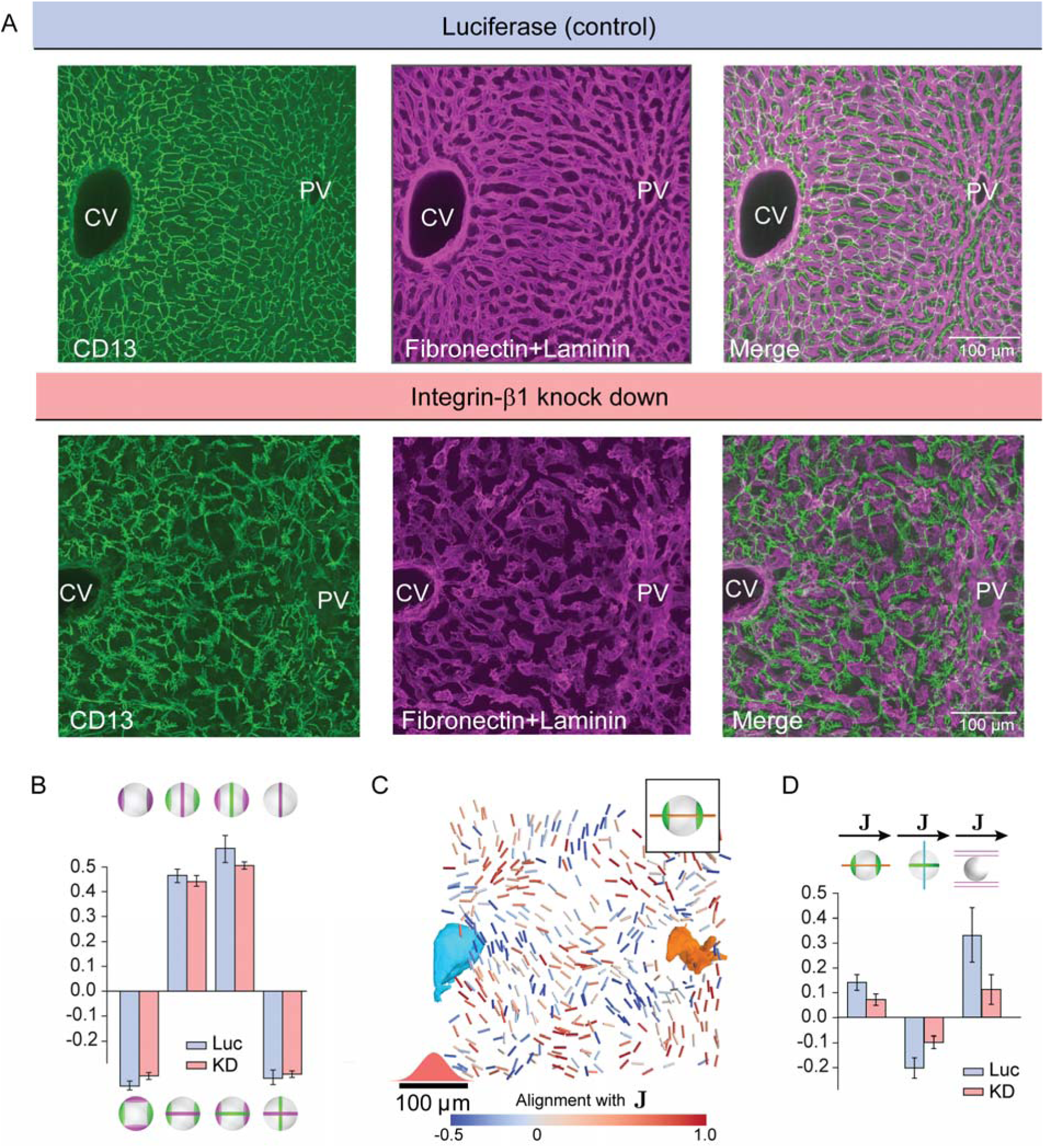
Liquid-crystal order, but not biaxial polarity of hepatocytes, is perturbed in integrin-β1 KD mice. **A)** Silencing integrin-β1 in the liver results in distortion of both bile canalicular and sinusoid networks, with reduced apparent alignment with the CV-PV axis in comparison to control conditions. Shown are representative samples for control conditions -top, siRNA against Luciferase (Luc) and integrin-β1 knock down -bottom, siRNA against integrin-β1 receptor (KD) stained for bile canalicular network (left, CD13 staining), sinusoid network (middle, fibronectin/laminin staining), and merge (right). All panels correspond to maximal intensity z-projection of 60*μm* of liver tissue. **B)** Individual hepatocytes retain their biaxial cell polarity in integrin-β1 KD, as revealed by cross-correlation analysis of nematic cell polarity axes for apical and basal plasma membrane patterns (analogous to Fig. 2F). **C, D)** In contrast, the alignment of biaxial cell polarity axes and the local preferred direction of the sinusoid network with **J** are reduced in integrin-β1 KD. Panel C shows bipolar cell polarity axes of apical plasma membrane color-coded according to their alignment with the local reference direction (**J**) (analogous to Fig. 3D). Panel D shows the quantification of alignment of apical bipolar axis, apical ring axis, and preferred sinusoid orientation with local reference direction (analogous to Fig. 3G). Statistics in B, C: mean+/−s.e. for n=5 animals (Luc) and n=4 animals (KD); statistical significance: panel B: *a*_1_ − *b*_1_: *p =* 0.015, *a*_1_ *− b*_2_:*p =* 0.277, *a*_2_ *− b*_1_: *p* = 0.118, *a*_2_ *− b*_2_*:p =* 0.410; panel *C*: *a*_1_ *−* **J**:*p* = 0.007, *a*_2_*−***J**:*p =* 0.002, *sinusoid −* **J**:*p*< 0.008, two-sided t-test assuming unequal variances).

## Discussion

Determining the structure of a protein, i.e. the three-dimensional arrangement of amino acids, allows making predictions on its function, intra- and inter-molecular interactions, as well as mechanisms of action and mutations that could alter its activity. Similarly, elucidating the structure of a tissue allows making predictions on how cells interact with each other and self-organize to form a functional tissue, including molecular mechanisms governing these processes (*23*). While some progress has been made in understanding 2D tissues (*13, 15, 24–29*) such as simple epithelia, the architecture of 3D tissues and its relation to function are poorly understood. The liver exemplifies this problem. Seventy years ago, Hans Elias pioneered an idealized structural model of liver tissue based on a crystalline order of cells (*6, 8*). Although his model captured some essential features of liver architecture, it could not explain the heterogeneity of cells and the amorphous appearance of the tissue.

In this study, we discovered novel design principles of liver tissue organization. We found that hepatocytes, BC and sinusoid networks are organized as a layered structure, with a spacing of about one hepatocyte diameter and orientation along the PV-CV axis, consistent with Elias’ model of hepatic plates. However, a breakthrough from our analysis was that, by using biaxial nematic tensors to describe hepatocyte polarity, we discovered that the polarity axes of individual hepatocytes are not random but display a liquid-crystal order on the scale of the lobule.

It has been proposed that the sinusoidal network forms a scaffold structure that guides hepatocyte polarity and BC network organization (*17, 18*). We propose an alternative organizational principle based on hierarchical levels of structural order (Fig. 5A). At the cellular level, hepatocytes display biaxial cell polarity of apical membrane distribution, distinct from the polarity in simple epithelia. At the multi-cellular level, the apical polarity axes of hepatocytes and the preferred direction of the sinusoidal network are aligned. Hepatocytes, BC and sinusoids exhibit a layered organization, where the layers are parallel to the veins. On the lobule level, we observed liquid-crystal order of hepatocyte polarity. This represents an intermediate state of order between highly ordered crystals and disordered liquids (Fig. 5B). The hierarchy of structural order could conceivably be explained by local rules of cell-cell communication in combination with global cues (e.g. morphogen gradients). Silencing Integrin-β1 provides a clue into the molecular mechanism underlying the local communication between hepatocytes and sinusoids. We found that the biaxial cell polarity of individual hepatocytes was maintained. In contrast, the liquid-crystal order was perturbed, i.e. tissue-scale alignment of cell polarity, and sinusoidal network anisotropy. Sinusoid-hepatocyte co-alignment could result from a self-organization mechanism, whereby sinusoids guide hepatocyte polarization, while hepatocytes provide instructive signals that guide sinusoidal network organization (Fig. 5C). This bidirectional communication provides a mechanism of self-organization for tissue development and maintenance. This points at a novel role of Integrin-β1 in orchestrating tissue structure by coupling the anisotropy of the sinusoidal network with the orientation of hepatocyte polarity, with boundary conditions set by the large veins.

**Fig. 5.**
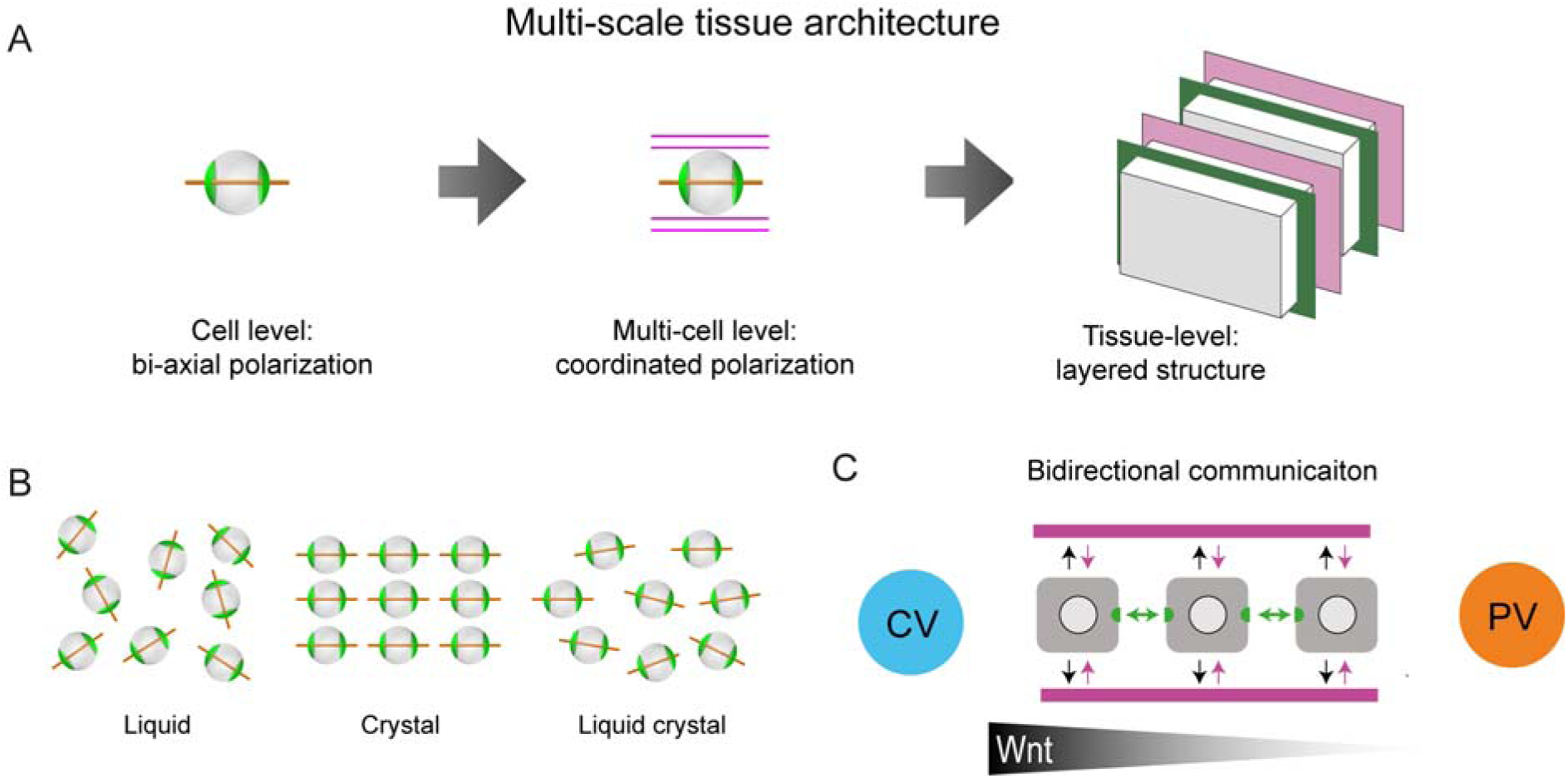
Proposed model of liver tissue architecture. **A)** Our work proposes a new multi-scale model of liver architecture, characterized by liquid-crystal order of hepatocytes with biaxial nematic cell polarity, co-alignment of hepatocyte polarity and preferred direction of the sinusoid network, and layered order of alternating sinusoid network and layers of hepatocytes with a thickness of one cell diameter. **B)** Cartoon representation of isotropic liquid with lack of positional and orientational order, crystalline order, and nematic liquid-crystal with orientational order. **C)** Schematic of bidirectional communication between sinusoids and hepatocytes. Long-range gradients, e.g. Wnt signaling, could provide alignment cues for the orientation biaxial cell polarity of hepatocytes (top-down organization), in addition to local interactions between hepatocytes and sinusoids (bottom-up organization).

In the liver, sinusoids and BC each form a single connected network. The two networks must be spatially separated, but also contact every hepatocyte to maximize the efficiency of fluid transport in the tissue. Furthermore, the networks are not tree-like, but highly interconnected and redundant with multiple loops and multiple contacts to each hepatocyte. This architecture confers robustness against local damage and failure. Our structural model of liquid-crystal order provides an explanation of how cells self-organize through local interactions to achieve this particular architecture, which is compatible with continuous homeostatic remodeling of the tissue.

Our model of liver tissue organization defines the road ahead. For example, are there intermediate (multi-cellular) units of structural organization between the cell and tissue level? Our study provides a general framework for elucidating the rules of cell-cell interactions and structural order of 3D tissues beyond the liver.

## Supporting information

## Acknowledgments

We thank Samuel Safran for stimulating discussions, and Stephan Grill, Peter Jansen, Carl Modes, Ivo Sbalzarini for a critical reading of the manuscript. We acknowledge generous allocation of computer time by the Center for Information Services and High Performance Computing (ZIH) at TU Dresden. We would like to thank the following Services and Facilities of the Max Planck Institute of Molecular Cell Biology and Genetics for their support: Biomedical Services (BMS) and Light Microscopy Facility (LMF).

## Author contributions

H.M-N, H.N, A.S, F.S-M, Y.K, F.J., B.M.F. and M.Z. conceived the project. H.N and F.S-M. performed the immunofluorescence experiments and imaging. H.M-N and Y.K. developed the image analysis algorithms. H.M-N and F.S-M. performed the 3D tissue reconstructions. H.M-N, A.S, W.deB. and L.B. performed the data analysis. A.S and B.M.F. developed the scheme for analysis of hepatocyte polarity and liquid crystal order. R.B and V.K provided tissue samples for KD experiments. H.M-N, A.S, F. S-M, K.M, L.B, Y.K, F.J, B.M.F. and M.Z. wrote the manuscript;

## Data and materials availability

All data available on request from the authors.

