## Supplementary material for "Liquid-crystal organization of liver tissue"

### Materials and Methods

#### Mice

To study liver tissue organization under normal conditions, 6 – 9 weeks old C57BL/6JOlaHsd mice (two males and one female) were purchased from Charles River Laboratory. For the knocked-down experiments, eight weeks old male mice were purchased from the same company. Integrin-β1 was knocked-down in mice (four males) by injecting siRNA formulated into lipidoid-based nanoparticles (LNP) that primarily target hepatocytes. An analogous treatment using siRNA against luciferase was used as control (five males). A complete description of the knock-down experiments can be found in (*1*). All procedures were performed in compliance with German animal welfare legislation and in pathogen-free conditions in the animal facility of the MPI-CBG, Dresden, Germany. Protocols were approved by the Institutional Animal Welfare Officer.

#### Sample collection and immunolabeling

Mice livers were fixed through intracardiac perfusion with 4% paraformaldehyde and post-fixed overnight at 4ºC with the same solution. Eight to ten serial slices were cut in a vibratome (thickness 100 μm), corresponding to a total thickness of 800-1000 μm. For immunostaining, we used anti-CD13 (Novus, cat NB100-64843, rat, 1/500), anti-fibronectin (Millipore, cat AB2033, rabbit, 1/1000), anti-laminin (Sigma, cat L9393, rabbit, 1/5000), anti-flk1 (R&D system, cat AF644, goat, 1/200), phalloidin-488 (LIFE technologies, cat A12379, 1/150) and DAPI (LIFE technologies, cat D1306, 1 μg/ml).

#### Immunostaining and optical clearing

Liver slices were permeabilized with 0.5% Triton X-100 in PBS for 60 minutes at room temperature (RT). Both, primary and secondary antibodies were diluted in TxBuffer (0.2% gelatin, 300 mM NaCl, 0.3% Triton X-100 in PBS) and each antibody was incubated with the tissue for 2 days at room temperature. For optical clearing we used a modified version of SeeDB (*2*). The first day the liver slices were incubated consecutively in 25% fructose for 4 hours, 50% fructose for 4 hours and 75% fructose overnight. The second day the samples were transferred to 100% fructose (100% wt/v fructose, 0.5% 1-thioglycerol, 0.1M phosphate buffer pH7.5) and the third day we left the samples in SeeDB solution (80.2% wt/wt fructose, 0.5% 1-thioglycerol, 0.1M phosphate buffer pH7.5) until the images were acquired at the microscope. Different concentrations of fructose were prepared diluting 100% fructose with water.

**Imaging and image analysis**

Liver samples were imaged in an upright multiphoton laser-scanning microscope (Zeiss LSM 780 NLO) equipped with Gallium arsenide phosphide (GaAsp) detectors. Liver slices were imaged twice at low (25x/0.8 Zeiss objective, 1 μm voxel size) and high resolution (63x/1.3 Zeiss objective, 0.3 μm voxel size), respectively. Low-resolution images were taken for the 3D reconstruction of big veins (central and portal veins) where the high resolution images were embedded. High-resolution images were acquired between selected central to portal vein (CV-PV) axes to resolve sub-cellular structures such as apical surfaces of hepatocytes. Both high- and low-resolution images were processed and segmented with the Motion Tracking software as described in (*3*).

#### Mollweide projection

We used the planar, pseudo-cylindrical Mollweide projection to visualize polarity patterns in 2D (Fig.2). The Mollweide projection preserves distances and areas at the expense of distorting shapes and is familiar from global maps of the world (*4*). For each cell, the apical plasma membrane domain of the cell surface was radially projected on a sphere placed at the volumetric center of the cell. To define a reference orientation for each cell, the bipolar axis of its projected basal plasma membrane ($b_1$) was chosen as the "north pole-south pole" axis for projection while the bipolar axis of the apical plasma membrane ($a_1$) defined the position of the zero meridian, which is placed vertically in the center of the projection. This relates the "poles" to the largest basal patches and the "equator" to apical patches with the largest apical patch pointing towards the reader.

#### Nematic tensors of polarity marker distribution of individual hepatocytes

In our software MotionTracking, the surface of a cell is represented as a triangulated mesh. For each vertex of the mesh, it is stored if this vertex was identified to belong to membrane patches rich in the apical polarity protein marker. Triangles are considered to belong to the apical plasma membrane patch if at least two vertices of the triangle have apical identity.

To compute the nematic tensor of apical polarity, triangles were first projected on a unit sphere to avoid distortions by non-spherical cell shapes, see Fig.2D. The center of the sphere is placed at the volumetric center of the cell. This projection assumes cell shapes to be star-convex with respect to the volumetric center of the cell. As a test, the sum of projected areas yielded $(4.035 \pm 0.086)\pi$, consistent with the surface area $4\pi$ of a unit sphere.

The nematic tensor of apical polarity is defined as a sum of all projected triangles with apical identify, indexed by $i \in I_{\text{apical}}$

(S3) $$N_{\alpha\beta} = \frac{3}{2}\frac{1}{\sum_i A_i}\sum_{i\in I_{\text{apical}}} A_i \left( n_\alpha^{(i)} n_\beta^{(i)} - \frac{1}{3}\delta_{\alpha\beta} \right)$$

Here, $A_i$ denotes the projected area of the projected triangle with index $i$ and $\mathbf{n}^{(i)}$ its surface normal vector. Einstein summation convention is assumed.

The rank-2 tensor $N_{\alpha\beta}$ is traceless and symmetric, and can thus be diagonalized with normalized eigenvectors $\mathbf{a}_1, \mathbf{a}_2, \mathbf{a}_3,$ and respective eigenvalues $\sigma_1$, $\sigma_2$, $\sigma_3$. Without loss of generality, we assume $\sigma_2 \leq \sigma_3 \leq \sigma_1$. We refer to $\mathbf{a}_1$as the bipolar axis, and $\mathbf{a}_2$ as the ring axis, and to $\sigma_1$, $\sigma_2$as their respective weights. Note that the eigenvectors are only determined up to sign, and thus each specify an axis. The third axis can be deduced from the other two as $\mathbf{a}_3 = \pm\mathbf{a}_1 \times \mathbf{a}_2$, while $\sigma_3 = -\sigma_1 - \sigma_2$. We did not observe strong correlations between these weights of polarity axes and the alignment of the axes with the lobule-level reference field $\mathbf{J}$ defined below (not shown).

The definition of the nematic tensor of basal polarity and the respective polarity axes is analogous.

Similarly, we define a preferred axis of the local sinusoid network. The skeleton of the network is characterized by segments with respective vectors $\mathbf{n}^{(i)}$and segment lengths $l^{(i)}$, for $i = 1, \ldots, N$. We define a nematic tensor that characterizes the anisotropy of the network

(S4) $$N_{\alpha\beta} = \frac{3}{2}\frac{1}{\sum_i l_i}\sum_{i=1}^{N} l_i \left( e_\alpha^{(i)} e_\beta^{(i)} - \frac{1}{3}\delta_{\alpha\beta}\right)$$

We refer to the axis parallel to the eigenvector corresponding to the largest eigenvalues of the sinusoid preferred axis.

**Local averaging of polarity axes**

For Fig.3B, D, E, and Fig.S7, local averaging of polarity axes was performed. Specifically, given unit vectors $\mathbf{a}^{(j)}$ that characterize a cell polarity axis of given type for individual hepatocytes (located at estimated cell center positions $\mathbf{x}^{(j)}$ labeled by an index $j$, we computed nematic tensors

$$\text{(S5)} \qquad \bar{N}_{\alpha\beta}(\mathbf{x}) = \tfrac{3}{2}\textstyle\sum_j w\left(\left|\mathbf{x}^{(j)} - \mathbf{x}\right|\right)\left( a_\alpha^{(j)} a_\beta^{(j)} - \tfrac{1}{3}\delta_{\alpha\beta}\right),$$

where $w(\mathbf{x})$ denotes a Gaussian kernel, with standard deviation chosen as 20 μm (approximately one hepatocyte diameter). In the respective plots, we displayed the principal axis with maximal eigenvalue of the weighted-mean tensors $\bar{N}_{\alpha\beta}(\mathbf{x}^{(i)})$ evaluated at the positions $\mathbf{x}^{(i)}$.

**Lobule-level reference system (J)**

The lobule-level reference system is defined by the location of the large vessels within the imaging volume by flux lines of solutions of the Poisson equation. The Poisson equation describes diffusive transport between spatially separated sources and sinks. Equivalently, the same solutions can be interpreted as an electrostatic potential, where positive and negative point charges correspond to the sources and sinks, respectively, and the electrostatic potential corresponds to a steady-state concentration field established by diffusion.

Below, we use the terminology of the electrostatic problem, which has the formal benefit that negative values of the $\chi$ field have a direct physical interpretation as a negative potential, whereas in the interpretation of concentration fields a homogeneous constant $\chi_0$ has to be added to the $\chi$ field to ensure that concentrations $c = \chi + \chi_0$ are non-negative.

The location of the large vessels (portal vein and central vein) are given as triangulated meshes as calculated by MotionTracking. Using the electrostatic analogy, point charges are placed at the location $\mathbf{r}_i$ of the triangle centers with point charges of strength $q_i$ proportional to the relative area of the triangle with respect to the total area of the corresponding vessel

$$\text{(S6)} \qquad q_i = \pm \frac{A_i}{\sum_j A_j}.$$

Here, the sum extends over all triangles of either the PV or CV mesh representation, respectively. The sign of the point charge $q_i$are opposite between the two vessel types, i.e. negative for portal vein and positive for central vein. This choice corresponds to a uniform charge surface density on the surfaces of the large vessels with total net charge equal to $\pm 1$, respectively.

A scalar field $\chi$ is calculated by superposition of the Green's functions of all the point charges on the veins

$$\text{(S7)} \qquad \chi = q_i \sum_i \frac{1}{|\mathbf{r}-\mathbf{r}_i|},$$

where the sum extends over all triangles of the PV and CV mesh representation. The value of this scalar field defines a positional value indicating location between PV and CV. Negative values indicate closeness to the portal vein, whereas positive values indicate closeness to the central vein. The gradient of this scalar field $\mathbf{J} = \nabla\chi$ gives a reference direction at all locations within the lobule. This field of reference directions is used to determine alignment of the nematic axes derived by the polarity of hepatocytes and the anisotropy of the sinusoid and BC networks.

**Nematic alignment parameter *S***

We consider an ensemble of axes, represented by unit vectors $\mathbf{e}^{(i)}$, together with an ensemble of reference axes with unit vectors $\mathbf{g}^{(i)}$, where $i = 1, \ldots, N$. We define the nematic alignment parameter as

(S8) $$S = \frac{1}{N}\sum_{i=1}^{N}\frac{3}{2}(\mathbf{e}^{(i)}\cdot\mathbf{g}^{(i)})^2 - \frac{1}{2}$$

We have $S = 1$for the case that each axis $\mathbf{e}^{(i)}$is perfectly parallel to its respective reference axis $\mathbf{g}^{(i)}$, whereas $S = -\frac{1}{2}$ for perfectly perpendicular relative alignment. For an isotropic orientation between axes and references axes, $S = 0$. We apply this general definition to cell polarity axes of hepatocytes and the preferred anisotropy axis of the local sinusoid network for $\mathbf{e}^{(i)}$, while the curvi-linear reference field $\mathbf{g}^{(i)} = \mathbf{J}$, evaluated at each cell position, provides local reference axes.

Our definition of the nematic alignment parameter S generalizes the known nematic order parameter (*5*, *6*). Specifically, if the reference axis in our definition is chosen constant and equal to the nematic symmetry axis of the ensemble, both definitions agree.

<u>Trivial baseline for nematic alignment parameter S for biaxial objects.</u>

We note a special property of the nematic alignment parameter for an ensemble of biaxial objects, characterized by respective unit vectors $\mathbf{e}_1^{(i)}, \mathbf{e}_2^{(i)}, \mathbf{e}_3^{(i)}$, $i = 1, \ldots, N$. Let $S_1, S_2, S_3$ be the respective nematic alignment parameters for each of the three different axes. Then, the maximum-likelihood estimate for $S_2$, given a known value for $S_1$, reads

(S9) $$S_2^{ML} = -\frac{1}{2}S_1$$

This relationship follows from $S_1 + S_2 + S_3 = 0$ and $S_2^{ML} = S_3^{ML}$. This trivial baseline is shown in Fig.3G as hatched bar for the correlation between the ring axis $\mathbf{a}_2$and $\mathbf{J}$.

**Cross-correlation analysis**

To investigate a possible layered order of liver tissue, we performed a cross-correlation analysis, see Fig.3H. We first selected a region of interest in the tissue sample, placed in the middle of the lobule. In this selected region, the field lines of the lobule-level reference field (**J**) are approximately straight and parallel, which facilitated analysis. The region is shown as a square in the inset of Fig.3H (enlarged in Fig.S6A). Inside the square region, the mean projected density of the sinusoid network (projected along the z-axis) based on segmented voxelated data is shown. Next, we investigated layered order between the positions of hepatocytes and the sinusoid and BC networks.

We calculated the cross-correlation between the mean projected density of the sinusoid network and an analogously defined mean projected density of hepatocytes (Fig.S6B). Specifically, we represent the mean projected density of the segmented sinusoids network by a 2D pixel array $\mathbf{S}[n,m]$and the averaged projected density of hepatocytes by a pixel array $\mathbf{H}[n,m]$, where $n$ and m are the indices of the pixels in the 2D images. The normalized cross-correlation between the respective images was then calculated as

$$C_{\mathbf{SH}}[k,l] := \frac{1}{N_{\mathbf{S}}+N_{\mathbf{H}}} \sum_{n,m} \frac{1}{\sigma_{\mathbf{S}}\sigma_{\mathbf{H}}} (\mathbf{S}[n,m]-\mu_{\mathbf{S}})(\mathbf{H}[n+k,m+l]-\mu_{\mathbf{H}}) \quad \text{(S10)}$$

where the sum runs over all pixels of $\mathbf{S}$. Here, $N_{\mathbf{S}}$denotes the total number of pixels of $\mathbf{S}$, $\mu_{\mathbf{S}} = \sum_{n,m} \mathbf{S}[n,m] / N_f$ is the average of $\mathbf{S}$, and $\sigma_{\mathbf{S}}$ the standard deviation, defined as $\sigma_{\mathbf{S}} = \sqrt{\sum_{n,m}\left(f[n,m]-\bar{f}\right)^2 / N_f}$. Analogous definitions apply to $N_{\mathbf{H}}$, $\mu_{\mathbf{H}}$, and $\sigma_{\mathbf{H}}$. Outside its valid range, the array $\mathbf{H}[n,m]$was zero-padded. The resultant 2D-cross correlation array $C_{\mathbf{SH}}[k,l]$ thus had twice the dimensions of $\mathbf{S}$.

We then computed the mean projection of this cross correlation array $C_{\mathbf{SH}}[k,l]$ on a line passing through $k = 0,\ l = 0$, and parallel to the blue reference line in the region of interest shown in the inset of Fig.3H. For this projection, we used a binning of 5 pixels, corresponding to $1.5\mu m$.

The use of a mean projection of 2D-cross correlation employed here instead of a conventional 1D-cross correlation along a single line reduced noise in the data and accounts for the fact of random phase shifts between neighboring layers.

An analogous cross correlation can be computed between the mean projected density of segmented hepatocytes and the BC network (Fig.S6C). The final projected cross correlations between sinusoid network and hepatocytes (magenta), as well as between sinusoid network and bile canaliculi network (green) are shown in Fig.3H (reproduced in Fig.S6D). As a control, Fig.S6E shows the analogous plot for a control direction perpendicular to the blue reference direction.).

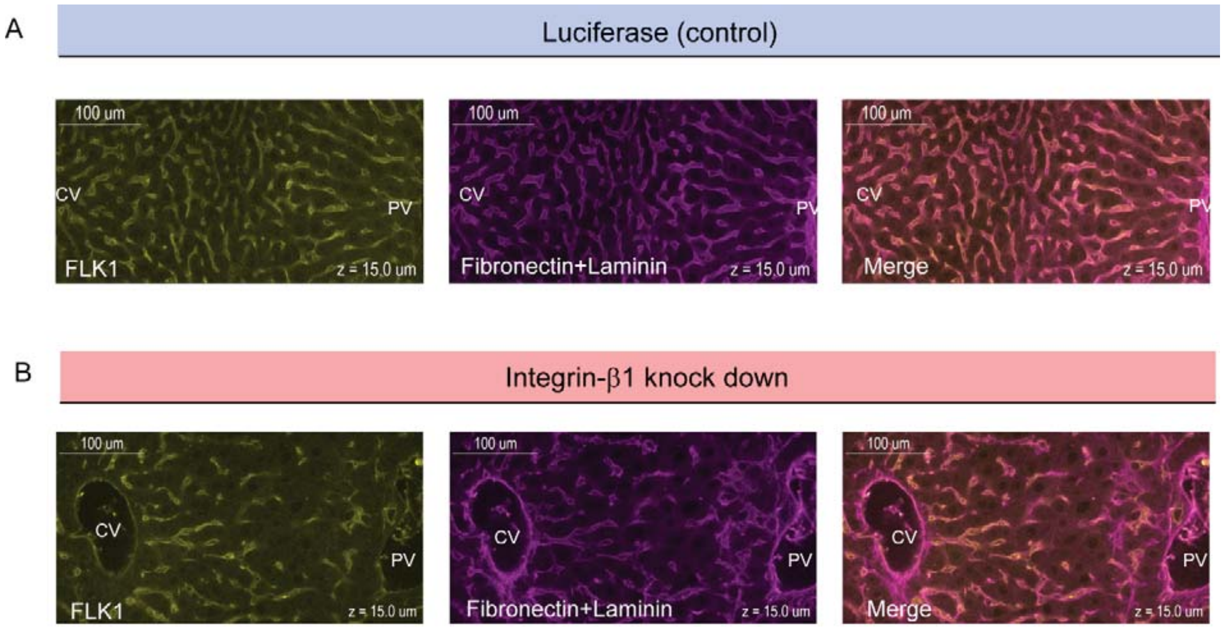


**Fig. S1. Colocalization of sinusoids and basal plasma membrane.**

High-resolution images of samples for control conditions (top, siRNA against Luciferase) and integrin-β1 knock down (bottom, siRNA against integrin-β1 receptor) are shown. Samples were stained for sinusoidal network (left, FLK1 staining) and extracellular matrix (middle, Fibronectin+Laminin staining). The right panels show images of overlapping staining. All panels correspond to a z-plane at 15 µm depth of liver tissue.

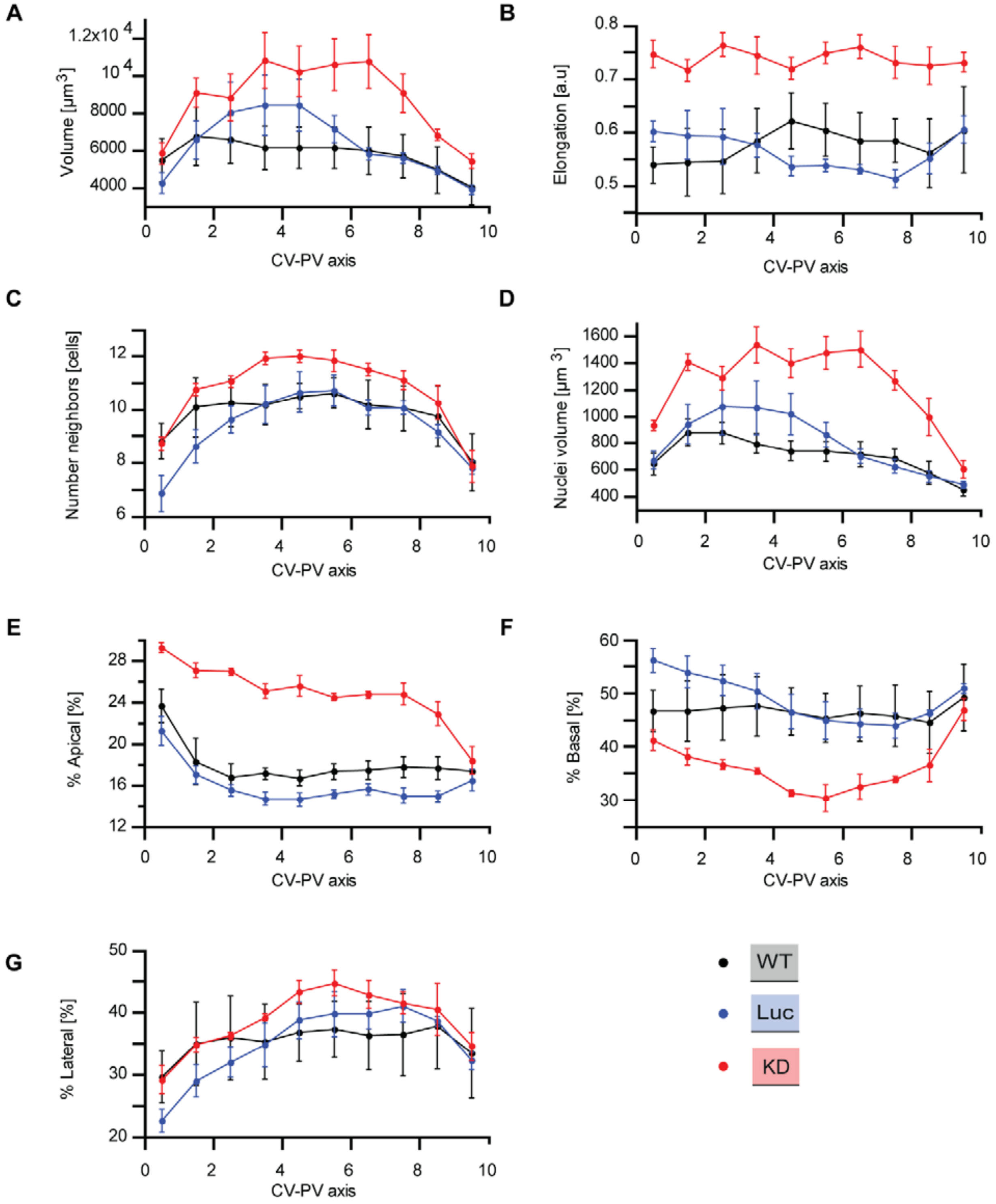


**Fig. S2. Quantitative structural parameters of hepatocytes along CV-PV axis.**

Structural parameters of hepatocytes for wild type (black), Luciferase control (blue), Integrin-β1 knockdown (red). As described in (3, 7) the CV-PV axis was divided in 10 zones (from 0 to 10) in order to show the variability of each parameter along this axis. **A)** Hepatocyte volume. **B)** Cell elongation. **C)** Number of neighboring hepatocytes per cell. **D)** Nuclear volume per cell. **E, F, G)** Percentage of apical, basal and lateral domains of the cytoplasmic membrane of individual hepatocytes, respectively. Statistics: n=3 independent samples from different animals (wild type), n=5 animals (Luciferase control), n=4 animals (Integrin-β1 knock-down). Error bars show s.e.m.

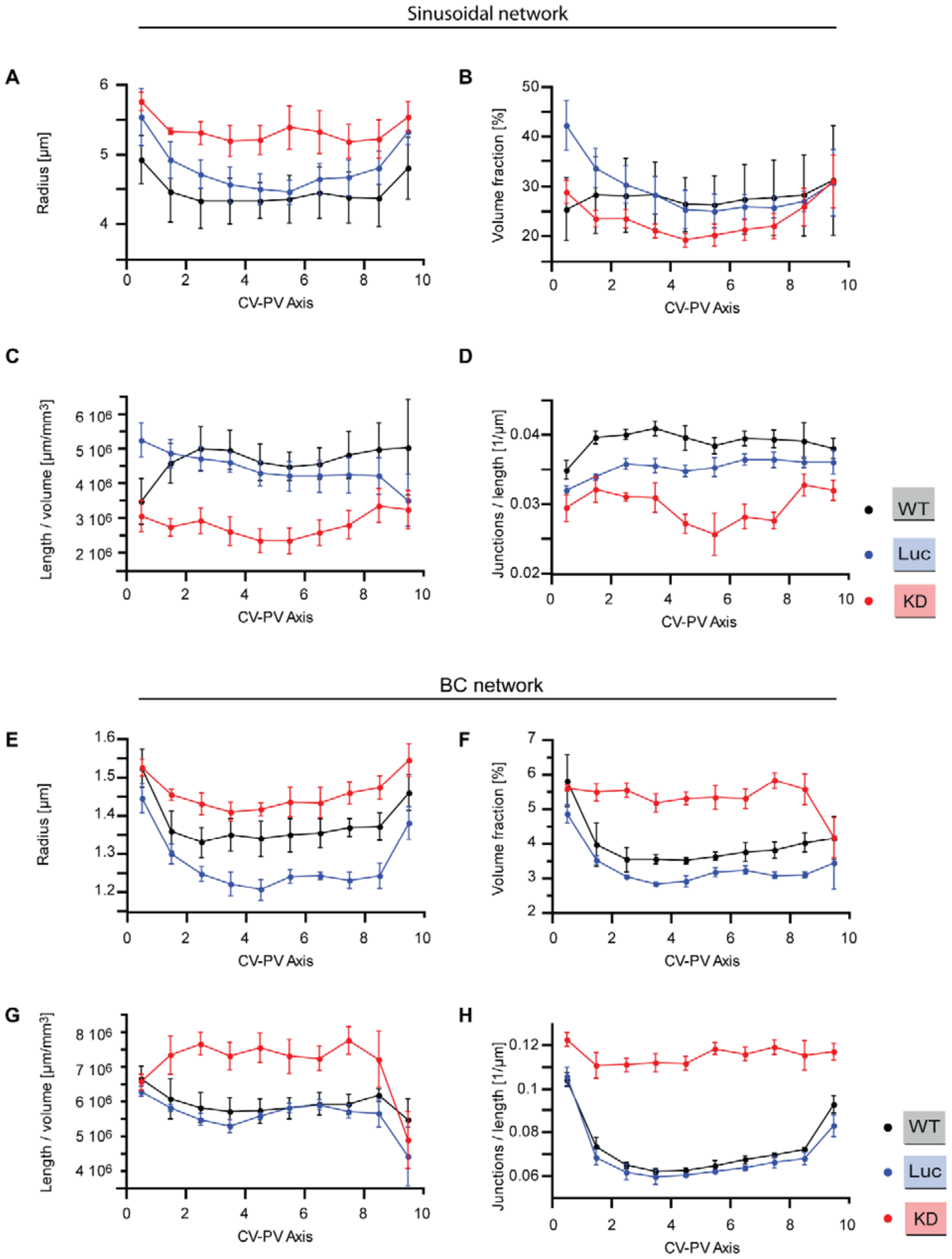
Sinusoidal network
A
Radius [µm]
CV-PV Axis
B
Volume fraction [%]
CV-PV Axis
C
Length / volume [µm/mm³]
CV-PV Axis
D
Junctions / length [1/µm]
CV-PV Axis
WT
Luc
KD
BC network
E
Radius [µm]
CV-PV Axis
F
Volume fraction [%]
CV-PV Axis
G
Length / volume [µm/mm³]
CV-PV Axis
H
Junctions / length [1/µm]
CV-PV Axis
WT
Luc
KD

**Fig. S3. Quantitative structural parameters of sinusoid and BC networks along CV-PV axis**

Structural parameters of the sinusoidal and BC for wild type (black), Luciferase control (blue), Integrin-β1 knockdown (red). The CV-PV axis was divided in 10 zones (from 0 to 10), showing change along the CV-PV axis. **A, E**) Estimated radius of the tubular networks. **B, F)** Percentage of tissue volume occupied by the network (excluding the volume occupied by CV and PV). **C, G)** Total length of the network per tissue volume unit. **D, H)** Number of branching points (junctions) per unit length. A junction is defined as a node of the network that has, at least, three branches. Statistics: n=3 independent samples from different animals (wild type), n=5 animals (Luciferase control), n=4 animals (Integrin-β1 knock-down). Error bars show s.e.m.

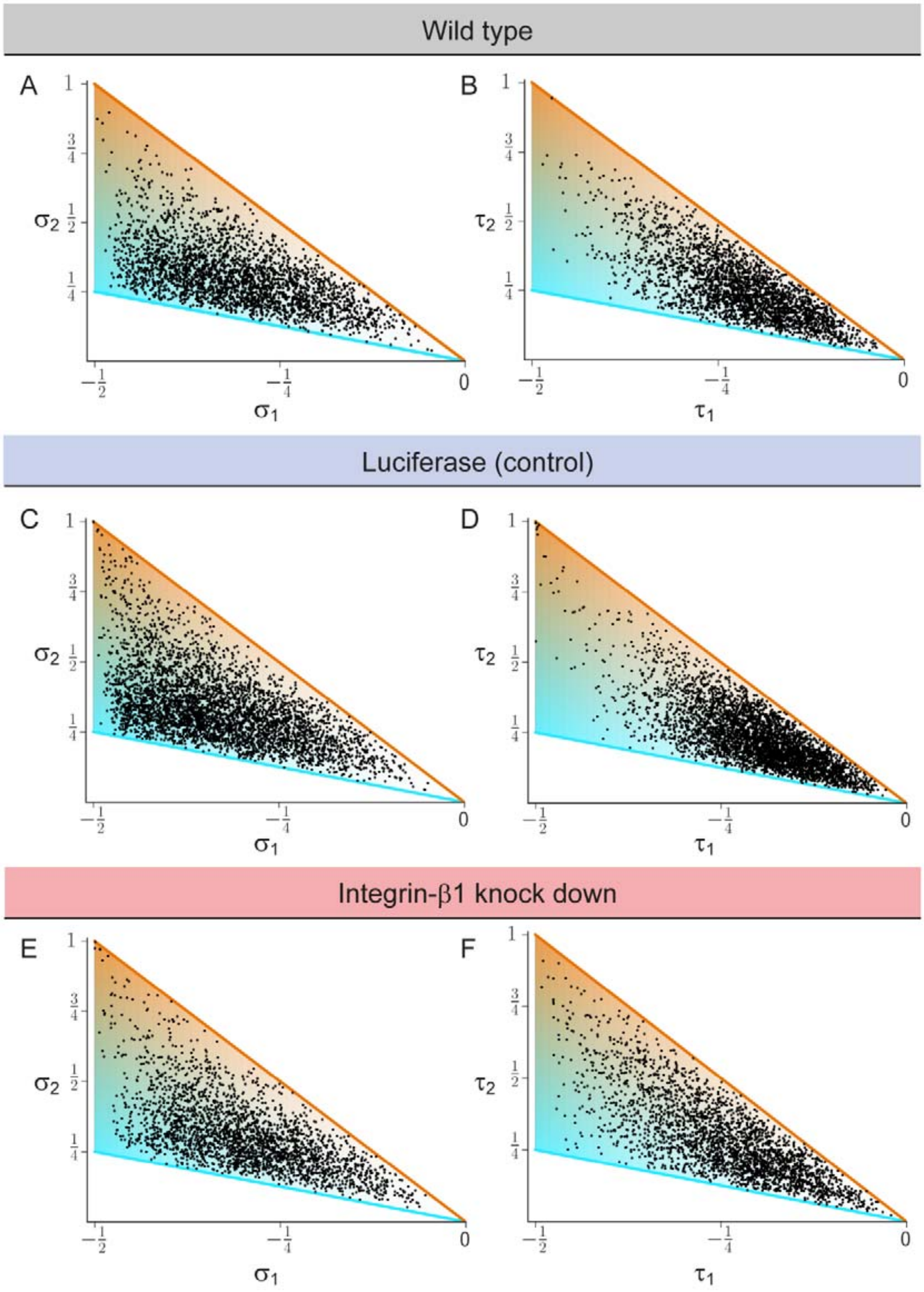


**Fig. S4. Biaxial cell polarity of basal plasma membrane distribution**

**A, C, E)** Respective weights $\sigma_1$ of bipolar axis ($a_1$) and $\sigma_2$ring axis ($a_2$) for reconstructed hepatocytes, defined in terms of the eigenvalues of the nematic cell polarity tensor for the apical plasma membrane distribution, for wildtype (panel A, identical to Fig.2E), Luciferase control (panel C), and Integrin-β1 knock-down (panel E). **B, D, F)**. Analogous scatter plots displaying the weight $\tau_1$ of the bipolar axis ($b_1$) and the weight $\tau_2$ of the ring axis ($b_2$) of basal plasma membrane distribution for reconstructed hepatocytes, defined in terms of the eigenvalues of the nematic cell polarity tensor for the basal plasma membrane distribution, for wildtype (panel B), Luciferase control (panel D), and Integrin-β1 knock-down (panel F). Data: n=3 independent samples from different animals (wild type), n=5 animals (Luciferase control), n=4 animals (Integrin-β1 knock-down).

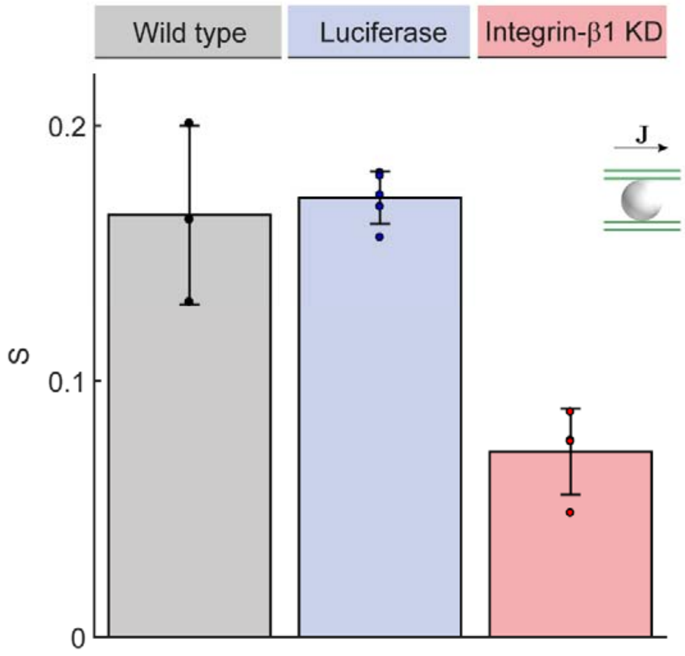


**Fig. S5. Anisotropy of BC network**

Quantification of alignment with local reference direction for the preferred direction of the local BC network surrounding each hepatocyte ($c_1$), analogous to Fig.3G. Statistics: n=3 independent samples from different animals (wild type), n=5 animals (Luciferase control), n=4 animals (Integrin-β1 knock-down). Error bars show s.e.m.

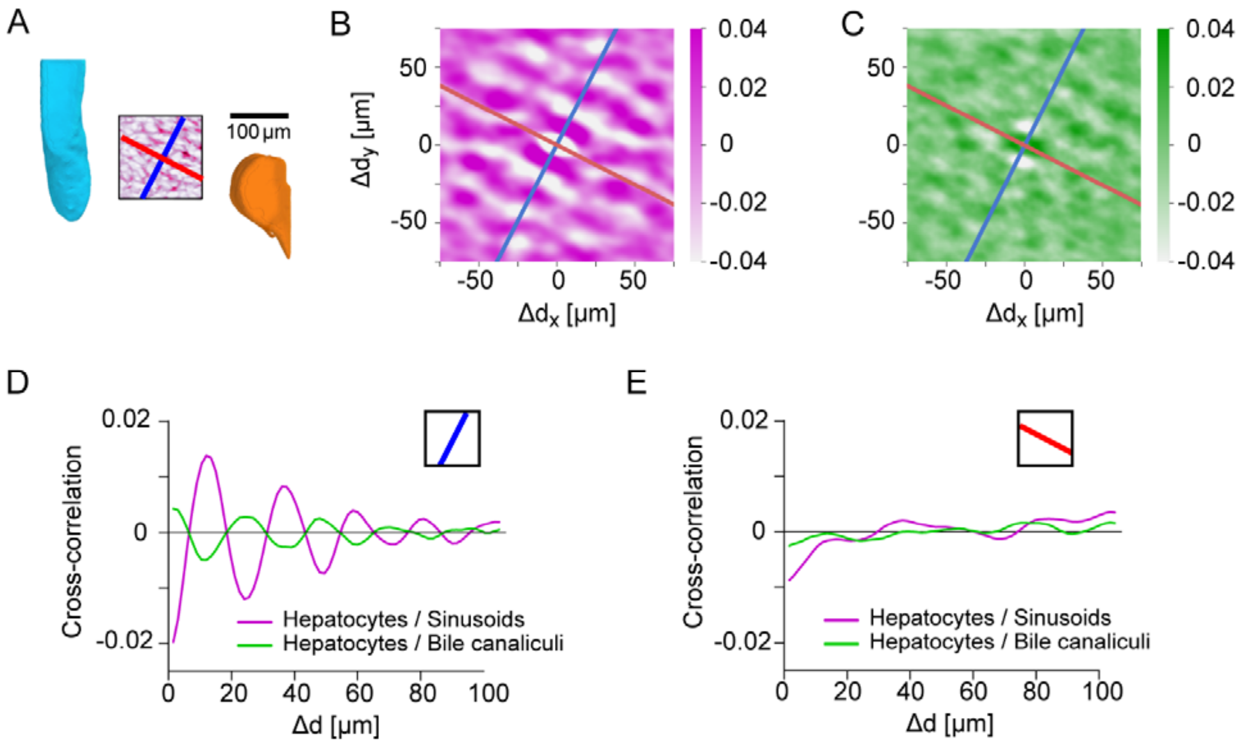


**Fig. S6. Control for layered order in the liver lobule**

**A)** Region-of-interest (ROI) in the central region of a liver lobule, with CV (cyan) and PV (orange) serving as landmarks; identical to inset of Fig.3H. Inside the ROI, the density of sinusoids is shown (average density projection along z-axis), together with a reference direction for layered order (blue) and a control direction (red), perpendicular to the reference direction. **B)** Two-dimensional cross-correlation between the projected density of hepatocytes and the projected density of sinusoids. **C)** Analogous to panel B for the cross-correlation between hepatocytes and the BC network. **D)** One-dimensional cross-correlations between the projected density of hepatocytes and sinusoids (magenta), as well as between hepatocytes and bile canaliculi (green), obtained from the two-dimensional cross-correlations by projection on the reference direction; identical to Fig.3H. **E)** Same as panel D, but for projection on the control direction. The absence of oscillatory signals is consistent with layered order of liver tissue with layers approximately orthogonal to the reference direction, but parallel to the control direction. Scale bar $100\mu m$.

252

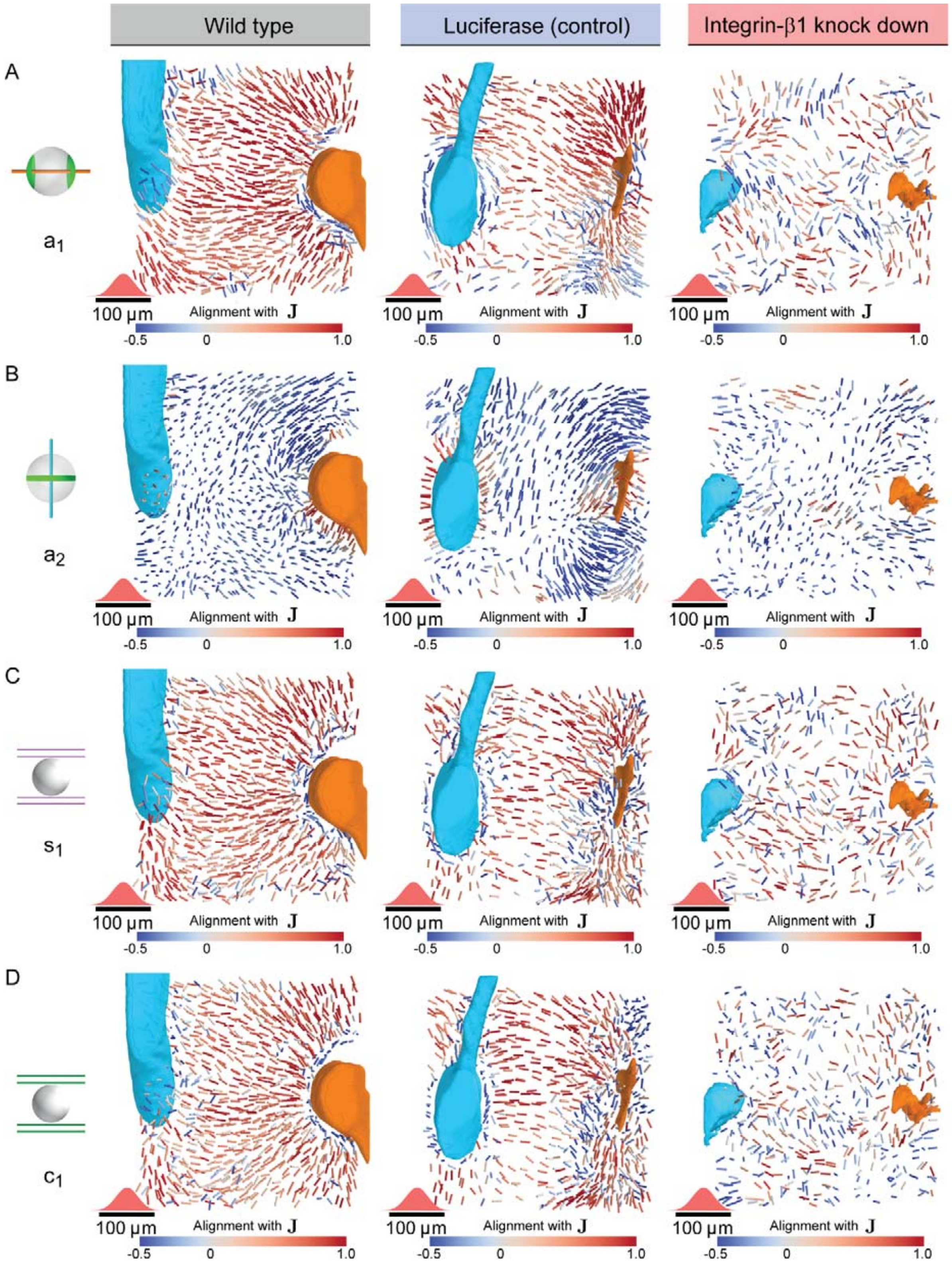

Wild type
Luciferase (control)
Integrin-β1 knock down
A
a1
B
a2
C
s1
D
c1
100 μm
Alignment with J
-0.5
0
1.0

**Fig. S7. Disturbed nematic liquid-crystal order in Integrin-β1 knock-down**

**A)** Bipolar cell polarity axes of apical plasma membrane distribution ($a_1$) shown as lines of constant length for individual hepatocytes at their respective position in the lobule after local averaging using a 3D Gaussian kernel at each hepatocyte cell center (standard deviation $20\mu m$, indicated in red above the scale bar) color-coded according to their alignment with the lobule-level reference field (**J**) for wildtype (left, identical to Fig.3D), Luciferase control (middle) and Integrin-β1 knock-down (right, identical to Fig.4C). **B)** Same as panel A, but for the ring axis of apical plasma membrane distribution ($a_2$), analogous to Fig.3E. **C)** Same as panel A, but for the preferred direction $s_1$ of the local sinusoid network surrounding each hepatocyte, analogous to Fig.3F. **D)** Same as panel A, but for the preferred direction $c_1$ of the local bile canaliculi network surrounding each hepatocyte.

**Movie S1. Supplementary movie for Fig.1C**

Digital reconstruction of large veins in mouse liver tissue, generated from low-resolution 3D images of serial slices (CV: cyan, PV: orange. Dimensions of imaging box: approximately 1mm × 1mm × 1mm.

**Movie S2. Supplementary movie for Fig.1E**

3D high-resolution reconstruction of main components of liver tissue: CV (cyan), PV (orange), sinusoidal network (magenta), bile canaliculi network (green), hepatocyte nuclei (random colors), and hepatocytes (random colors). Dimensions of imaging box: approximately $400\mu m$ × 400µm × 100µm.

**Movie S3. Supplementary movie for Fig.1F**

Single hepatocyte showing apical (green), basal (magenta) and lateral (grey) plasma membrane domains, reconstructed from high-resolution 3D images.
